# NuMA deficiency causes micronuclei via checkpoint-insensitive k-fiber minus-end detachment from mitotic spindle poles

**DOI:** 10.1101/2022.10.04.510904

**Authors:** Marvin van Toorn, Amy Gooch, Susan Boerner, Tomomi Kiyomitsu

## Abstract

Micronuclei resulting from improper chromosome segregation foster chromosomal instability in somatic cell division cycles. To prevent micronuclei formation, bundled kinetochore-microtubules called k-fibers must be properly connected to all sister kinetochores on chromosomes via their plus-ends, whereas k-fiber minus-ends must be clustered at the two opposing spindle poles throughout mitosis. The bipolar attachment between sister kinetochores and k-fiber plus-ends is carefully monitored by the spindle assembly checkpoint and further promoted by error-correction mechanisms. However, how k-fiber minus-end clustering is maintained and monitored remains poorly understood. Here, we show that degradation of the Nuclear Mitotic Apparatus (NuMA) protein by auxin-inducible degron technologies in human cells results in micronuclei formation through k-fiber minus-end detachment from focused spindle poles during metaphase. Importantly, this k-fiber minus-end detachment creates misaligned chromosomes that maintain chromosome biorientation and do not activate the mitotic checkpoint, resulting in lagging chromosomes in anaphase. Moreover, we find that NuMA depletion causes centrosome clustering defects in tetraploid cells, leading to an increased frequency of multipolar divisions. Together, our data indicate that NuMA-mediated minus-end clustering of k-fibers and spindle microtubules is critical for faithful chromosome segregation. Similar to erroneous merotelic kinetochore attachments, detachment of k-fiber minus-ends from metaphase spindle poles evades spindle checkpoint surveillance and may therefore be a source of genomic instability in dividing cells.

## Introduction

During eukaryotic cell division, a microtubule-based bipolar spindle is assembled to equally distribute duplicated chromosomes to daughter cells^1^. Dysfunction of the mitotic spindle can lead to abnormal chromosome segregation followed by aneuploidy and micronuclei formation, which further promote extensive chromosomal instability^2,3^. During mitotic spindle formation, many short microtubules are locally nucleated around chromatin, and from centrosomes and pre-existing microtubules^1,4^. These microtubules are subsequently elongated, bundled, crosslinked, transported, or captured by microtubule-binding proteins or motors, and coordinately organized in a large bipolar spindle^1^.

For successful chromosome segregation, the mitotic spindle must fulfill at least two requirements. First, the dynamic plus-ends of bundled kinetochore-microtubules (k-fibers) must be properly attached to all chromosomes via kinetochores^5^. Especially, k-fibers captured by sister kinetochores must be linked to opposite spindle poles to establish chromosome biorientation before anaphase onset. Second, the less dynamic and often γ-tubulin-capped minus-ends of microtubules^6^ must be properly focused at the two opposing spindle poles. In particular, k-fiber minus-end clustering must be established during metaphase and maintained during anaphase^7^. In most cases, focused spindle poles are located near centrosomes in animal mitosis, and detachment of k-fiber minus-ends from the focused poles would become a potential risk for abnormal segregation.

Multiple mechanisms promote the establishment and maintenance of chromosome biorientation in early mitosis. First, the spindle assembly checkpoint (SAC) proteins accumulate on unattached kinetochores after nuclear envelope breakdown (NEBD) to monitor kinetochore-microtubule attachment state during chromosome congression^8^ and delay anaphase onset until all chromosomes are properly attached to microtubules from opposing spindle poles^9^. In addition, error correction mechanisms sense and correct abnormal kinetochore-microtubule attachments^10^. However, in contrast to these plus-end monitoring processes, whether and how k-fiber minus-end focusing is monitored remains poorly understood.

The Nuclear Mitotic Apparatus (NuMA) protein recognizes microtubule minus-ends and clusters them around spindle poles in cooperation with dynein and dynactin in human cells^11,12^. NuMA dysfunction causes spindle-pole focusing defects in early mitosis and lagging chromosomes in anaphase^11,13–16^. Although NuMA depletion has been reported to cause micronuclei independently of its mitotic spindle functions by promoting nuclear dynamics and mechanics^17^, how spindle-pole focusing defects lead to abnormal chromosome segregation remains largely unclear. In this study, we analyzed the molecular mechanisms and consequences of spindle-pole focusing defects in NuMA-depleted human cells. We found that NuMA depletion causes chromosome mis-alignment due to k-fiber minus-end detachment from the focused spindle poles in metaphase, which is not sensed or corrected by the SAC and leads to abnormal chromosome segregation.

## Results and Discussion

### NuMA depletion causes micronuclei formation due to k-fiber minus-end detachment from metaphase spindle poles

NuMA depletion by auxin-inducible degron (AID) tagging causes spindle-pole focusing defects in human HCT116 cells^15,16^. To understand the consequence of these spindle-pole focusing defects, we first analyzed interphase phenotypes 24 hours after addition of doxycycline (Dox) and auxin (IAA), which induce degradation of endogenous mAID-tagged NuMA (NuMA-mAID-mClover-FLAG, hereafter described as NuMA-mACF)^15,18^. Remarkably, ~40% of NuMA-depleted cells contained one or more micronuclei, whereas micronuclei were hardly detected in control cells (Figure 1A and 1B). Similar results were obtained when we depleted endogenous NuMA using the recently developed AID2 system^19^ (Figure S1A-C, where endogenous NuMA was tagged with mAID-mCherry), which overcomes leaky degradation of mAID-fusion proteins and more efficiently and uniformly degrades the target protein with lower amounts of auxin analogue, 5Ph-IAA.

**Figure 1.**
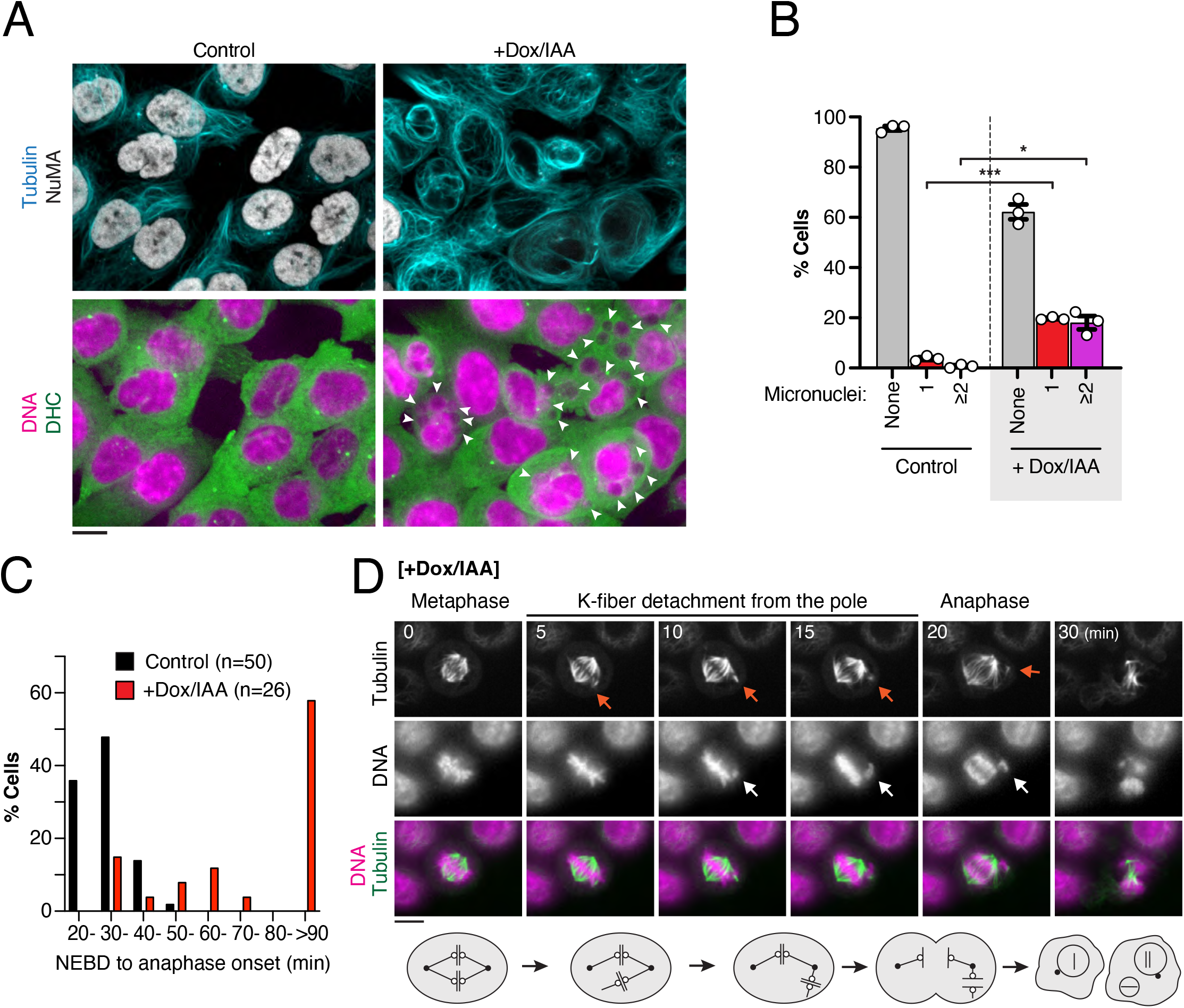
NuMA depletion causes micronuclei formation due to k-fiber minus-end detachment from focused spindle poles during metaphase. **(A)** Representative live-cell fluorescent images showing the presence of micronuclei in interphase NuMA-mACF knock-in cells. Cytoplasmic dynein was visualized by the DHC subunit to discern micronuclei^15^. Where indicated, endogenous NuMA was degraded by addition of 2 μg/mL doxycycline (Dox) and 500 μM IAA 20 hr prior to imaging. Arrowheads indicate micronuclei. **(B)** Quantification of micronuclei formation in control and NuMA-depleted interphase NuMA-mACF KI cells. Plotted values are averages of n = 3 independent experiments ± SEM. **(C)** Quantification of time elapsed from NEBD until anaphase onset in control and NuMA-depleted cells after release from double thymidine block. ≥90 min indicates cells that did not enter anaphase within 90 min during 3 hr time-lapse imaging. **(D)** Top: Live-cell fluorescent images of NuMA-depleted cells showing k-fiber minus-end detachment (red arrows) and chromosome misalignment (white arrows) from the focused spindle pole, followed by anaphase entry. Bottom: Schematic representation of the k-fiber minus-end detachment phenotype. Scale bars = 10 μm.

To study whether and how micronuclei are created during mitosis, we performed live-cell imaging of mitotic NuMA-depleted cells. Under our conditions (see Methods for details), control cells entered anaphase about 30 min after NEBD (31 ± 6.6 min, n = 50) (Figure 1C). Chromosomes aligned to form the metaphase plate and were properly segregated to daughter cells (Figure S1D). In contrast, 57.7% of NuMA-depleted cells showed mitotic delay (>90 min, n = 15) with chromosome congression defects (Figure 1C and Figure S1E Cell #1), consistent with a previous study^20^. The remaining 42.3% of cells entered anaphase with normal timing (32.5 ± 2.5 min, n = 4) or slight mitotic delay (57.1 ± 10.6 min, n = 7) (Figure 1C and 1D, Figure S1E Cell #2). In the latter case, cells showed transient delay in prometaphase, but finally formed a bipolar spindle with properly aligned chromosomes (Figure 1D, t = 0). However, some k-fibers were subsequently detached from the focused spindle poles, resulting in chromosome misalignment (Figure 1D, t = 5 - 15). In all cases observed, the minus end of the k-fibers dissociated from only one pole, resulting in the dissociated microtubules and associated chromosome moving in an arc toward the opposing, still clustered, pole (Figure 1D). In certain cases, this phenotype was even observed for multiple distinct k-fibers in the same cell (Figure S1E Cell #2 t = 40, also see Figure S2F, t =15 and 3A, t = 33). Importantly, these cells entered anaphase despite the presence of the misaligned chromosomes (Figure 1D, t = 20). Other cells displayed similar k-fiber detachment just before or during anaphase, which led to lagging chromosomes followed by micronuclei formation (Figure S1E Cell #2). These results suggest that NuMA depletion causes micronuclei in part due to k-fiber minus-end detachment from focused spindle poles during metaphase or anaphase.

### Mad2 does not accumulate on misaligned chromosomes generated by k-fiber minus-end detachment from spindle poles

SAC proteins accumulate on unattached kinetochores and delay anaphase onset until all chromosomes are properly captured by microtubules^9^. To understand whether the misaligned chromosomes generated by NuMA depletion are similarly sensed by checkpoint proteins, we next sought to visualize a key checkpoint protein Mad2^8^ together with the kinetochore protein Mis12^21^ upon NuMA depletion using the AID2 system. We established quadruple knock-in cell lines (Figure S2A) and subsequently analyzed the localization of endogenous Mis12 and Mad2 in living cells (Figure 2A). To deplete NuMA while minimizing the duration before mitotic entry, we induced NuMA degradation in cells arrested in G2 with the CDK1 inhibitor RO-3306^22^ and analyzed NuMA-depleted mitotic phenotypes immediately after RO-3306 washout (Figure S2B, see Methods for details). Similar to NuMA-depletion by the classical AID technology (Figure 1C and D), NuMA-depletion via the AID2 system resulted in ~35% of cells displaying k-fiber minus-end detachment from spindle poles, resulting in chromosome misalignment in metaphase (Figure 2A, t = 12 - 15, and Figure S2C) and lagging chromosomes in anaphase (Figure 2A, t = 18).

**Figure 2.**
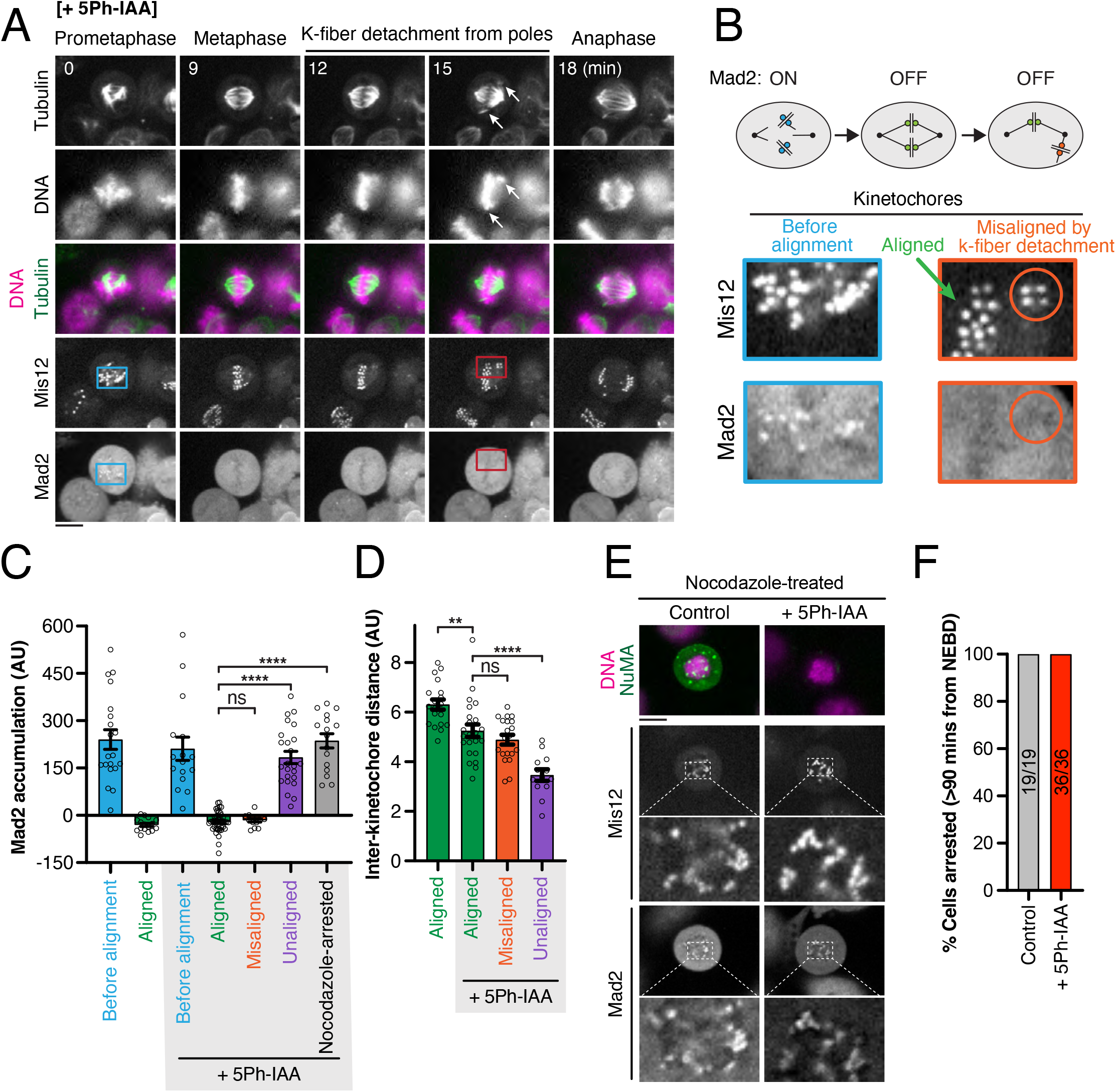
Misaligned chromosomes generated by k-fiver minus-end detachments from spindle poles are not sensed by the spindle assembly checkpoint effector Mad2. **(A)** Representative live-cell fluorescent images showing k-fiber minus-end detachment and anaphase entry in NuMA-depleted OsTIR1 (F74G) / NuMA-mAID-mCherry / MIS12-SNAP / Mad2-mClover quadruple KI cells released from RO-3306-induced G2/M arrest. **(B)** Top: Schematic representation of Mad2 localization at kinetochores before, during and after losing alignment on the metaphase plate. Bottom: Enlargement of indicated areas in (A), showing Mad2-positive kinetochores before alignment (sky blue square) and Mad2-negative kinetochores after k-fiber minus-end detachment during metaphase (orange circle). **(C)** Quantification of Mad2 accumulation on the indicated categories of kinetochores in control and NuMA-depleted KI cells shown in (A). Plotted values are averages ± SEM from n ≥ 13 kinetochores imaged in 3 independent experiments. **(D)** Quantification of inter-kinetochore distance on the indicated categories of sister kinetochore pairs. Plotted values are averages ± SEM from n ≥ 13 sister kinetochore pairs imaged in 3 independent experiments. **(E)** Fluorescent images of control and NuMA-depleted KI cells treated with 330 nM nocodazole for 4 hr after release from RO-3306-induced G2/M arrest. Insets show accumulation of Mad2 on kinetochores, marked by Mis12, in both control and NuMA-depleted cells. Scale bars = 10 μm. **(F)** Quantification of nocodazole-induced mitotic arrest in control and NuMA-depleted KI cells, which was defined as ≥90 min from NEBD. Counted cells were imaged in 3 independent experiments.

As expected, Mad2 accumulated on kinetochores before alignment in prometaphase and dissociated from aligned kinetochores during metaphase in both control and NuMA-depleted cells (Figure 2A-C and Figure S2D-G). Importantly, Mad2 was not detected on the misaligned kinetochores generated by k-fiber minus-end detachment in NuMA depleted cells (Figure 2A-C and Figure S2F-G). In all cases observed (n = 37), Mad2 was not targeted to these misaligned kinetochores after k-fiber minus-end detachment, and cells entered anaphase with lagging chromosomes (Figure 2A, t = 18, and Figure S2F, t = 42). We next measured inter-kinetochore distances of the misaligned chromosomes to understand whether sister kinetochores are still properly pulled on by microtubules after k-fiber minus-end detachment. Inter-kinetochore distance was also similar between aligned and misaligned kinetochores in NuMA-depleted cells (Figure 2D), although inter-kinetochore distance on aligned kinetochores was slightly reduced compared to that of the control, consistent with previous studies^13,14^. In contrast, Mad2 was detected on unaligned kinetochores which showed congression failures during prometaphase and never properly aligned on the metaphase plate during observation (Figure 2C and Figure S2H-I). In these Mad2-positive unaligned kinetochores, inter-kinetochore distance was reduced compared to Mad2-negative aligned and misaligned chromosomes (Figure 2D), suggesting that the Mad2-negative misaligned kinetochores are still captured and physically pulled on by microtubules. Mad2 also accumulated on kinetochores when microtubule polymerization was inhibited by nocodazole treatment (Figure 2C and 2E), which consequently caused mitotic arrest in both control and NuMA-depleted cells (Figure 2F). These results indicate that the SAC can be normally activated in the absence of NuMA, but that misaligned chromosomes generated by k-fiber minus-end detachment from metaphase spindle poles still maintain chromosome biorientation, and thus are not sensed by Mad2.

### BubR1 shows no accumulation on misaligned chromosomes caused by k-fiber minus-end detachment from spindle poles

Mad2 is targeted to unattached kinetochores, where it forms a potent anaphase inhibitor, mitotic checkpoint complex (Mad2-Cdc20-BubR1-Bub3)^9^. Since BubR1-Bub3 is targeted to unattached kinetochores independently of Mad2^9^, we next sought to analyze the localization of BubR1 in NuMA-depleted cells (Figure S3A). BubR1 behaved similarly to Mad2 in that it accumulated on most prometaphase kinetochores before alignment and dissociated from aligned kinetochores (Figure 3A-C). However, unlike Mad2, low BubR1 signals were still detected on aligned metaphase kinetochores in both control and NuMA-depleted cells (Figure 3B and 3C). BubR1 levels on misaligned kinetochores were equivalent to those on aligned kinetochores (Figure 3B and C). On the other hand, BubR1 was significantly enriched on unaligned kinetochores with shorter inter-kinetochore distance (Figure 3C-D and Figure S3E-F) and on kinetochores in nocodazole-arrested cells (Figure 3E and 3F). These results support our conclusion that misaligned kinetochores generated by k-fiber minus-end detachment maintain bipolar attachment and thus do not activate mitotic checkpoint signaling.

**Figure 3.**
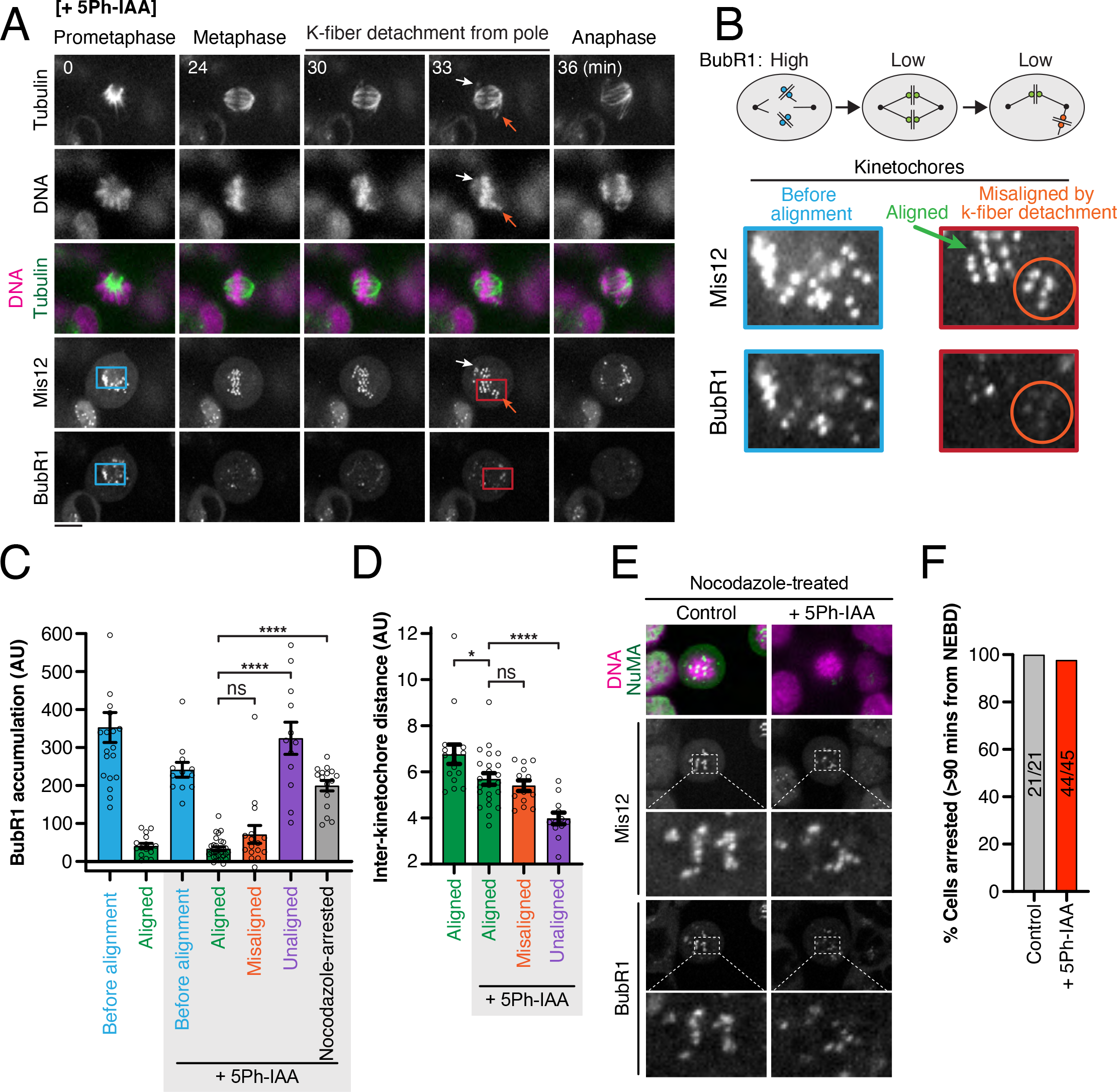
BubR1 does not accumulate on misaligned kinetochores caused by k-fiber minus-end detachment during metaphase. **(A)** Representative live-cell fluorescent images showing kMT detachment and anaphase entry in NuMA-depleted OsTIR1(F74G) / NuMA-mAID2-mCherry / MIS12-SNAP / BubR1-mClover quadruple KI cells released from RO-3306-induced G2/M arrest. Orange arrow (t = 33) indicates detachment of two k-fibers from the metaphase spindle pole. **(B)** Top: Schematic representation of BubR1 accumulation at kinetochores before, during and after losing alignment to the metaphase plate. Bottom: Enlargement of indicated areas in (A), showing BubR1-high kinetochores before alignment (e.g. during prometaphase) and BubR1-low kinetochores after k-fiber minus-end detachment from the focused spindle pole during metaphase. **(C)** Quantification of BubR1 accumulation on the indicated categories of kinetochores in control and NuMA-depleted KI cells. Plotted values are averages ± SEM from n ≥ 12 kinetochores imaged in 3 independent experiments. **(D)** Quantification of inter-kinetochore distance on the indicated categories of sister kinetochore pairs. Plotted values are averages ± SEM from n ≥ 12 sister kinetochore pairs imaged in 3 independent experiments. **(E)** Fluorescent images of control and NuMA-depleted KI cells treated with 330 nM nocodazole for 4 hr after release from RO-3306-induced G2/M arrest. Insets show BubR1 accumulation on kinetochores in both control and NuMA-depleted cells. **(F)** Quantification of nocodazole-induced mitotic arrest in control and NuMA-depleted KI cells from 3 independent experiments. Scale bars = 10 μm.

### NuMA depletion causes centrosome clustering defects in tetraploid cells

When we analyzed NuMA-depletion phenotypes using a knock-in clone expressing BubR1-mClover, we unexpectedly found an increase in multipolar division (Figure S3B and S3G). Since these cells specifically had a large chromosome mass, we hypothesized that NuMA might be required to maintain not only k-fiber clustering in diploid cells, but also centrosome clustering in tetraploid cells by crosslinking spindle microtubules emanating from neighboring centrosomes. To test this hypothesis, we induced tetraploidy by inhibiting cytokinesis using the actin polymerization inhibitor cytochalasin D (Figure S4A, see Methods for details). Under our conditions, bipolar spindles were observed in ~70% of tetraploid-like control cells. However, in NuMA-depleted cells, the frequency was reduced to ~40% and multi-polar spindles were predominantly observed (Figure S4B and S4C: we got similar results using two different knock-in clones, Mad2- or BubR1-mClover expressing cells).

To further confirm these results, we next visualized centrosomes by creating knock-in cell lines expressing γ-tubulin-SNAP (a centrosome marker, Figure S4D) and analyzed metaphase cells with >3 large, visually discernable, γ-tubulin signals after the induction of tetraploidy. Under our conditions, 50-60% of control cells assembled a bipolar spindle by clustering two large γ-tubulin signals at each pole (Figure 4A and 4B). However, such centrosome-clustering was observed in only 20-30% of NuMA-depleted cells, and instead, non-bipolar spindles with more than 3 γ-tubulin signals were predominantly observed in NuMA-depleted cells (Figure 4A and 4B: the results were confirmed with two different γ-tubulin-SNAP knock-in clones, #1 and #2). These results suggest that NuMA crosslinks kinetochore- or spindle-microtubules connected to neighboring centrosomes and promotes bipolar division by centrosome clustering in tetraploid cells.

**Figure 4.**
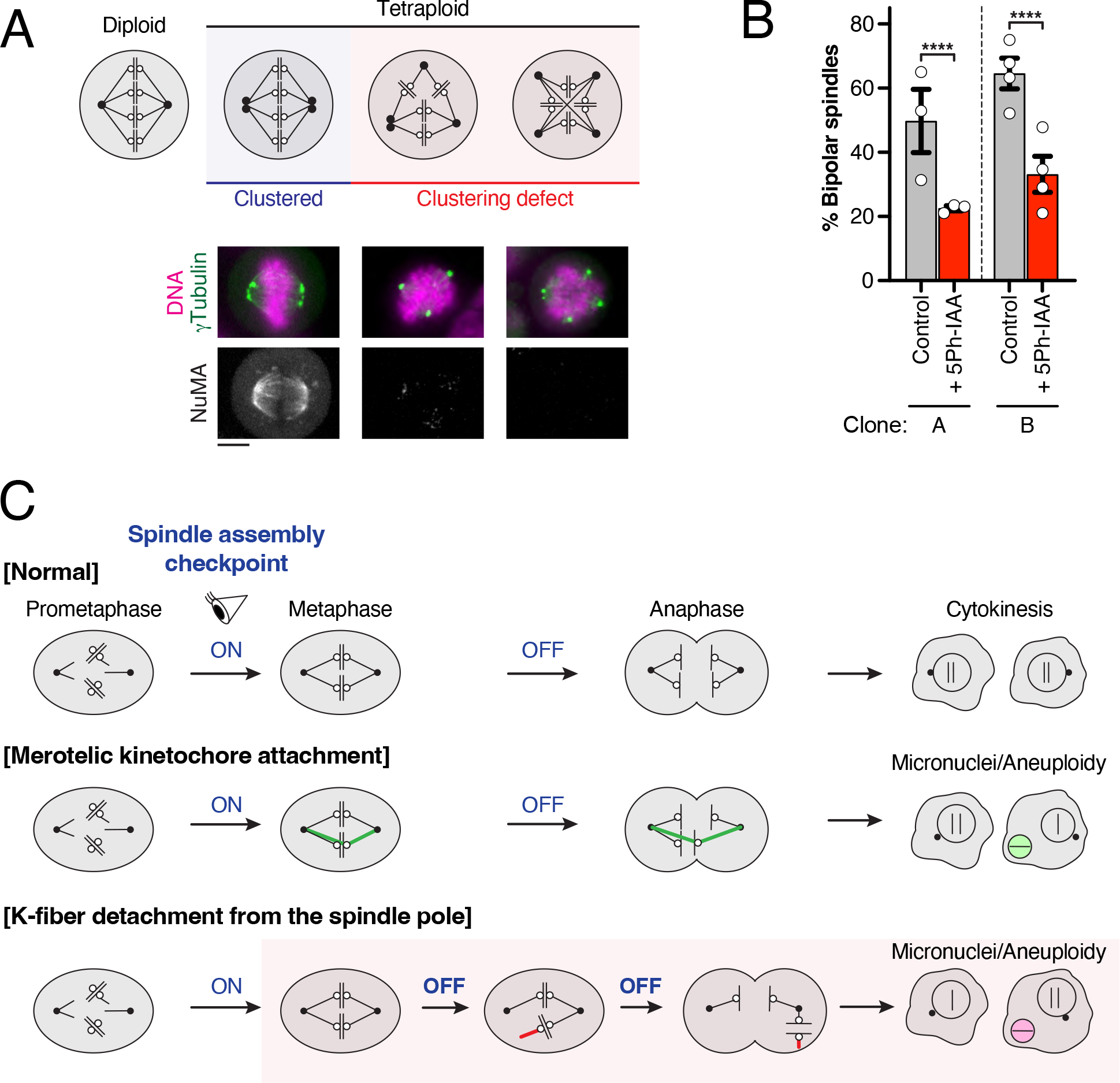
NuMA is required for efficient centrosome clustering in tetraploid cells. **(A)** Top: Schematic representation of proper (blue, left) and improper (red, right) centrosome (black dots) clustering in tetraploid cells. Bottom: Representative fluorescent images of control and NuMA-depleted triple KI cells showing proper and improper clustering of centrosomes, marked by γ-Tubulin protein TUBG1. **(B)** Quantification of the percentage of bipolar spindles in tetraploid metaphase or early anaphase cells with >3 clear distinguishable centrosomes. Plotted values are average percentages ± SEM from n ≥ 3 independent experiments, in which >17 cells were counted each time. **(C)** Model showing micronuclei formation due to SAC-insensitive abnormal chromosome segregation pathways. In contrast to the normal situation (top panels), erroneous merotelic attachment (shown in green in the middle panels) is not sensed by the SAC and causes micronuclei. Although k-fiber minus-end detachment from the metaphase spindle pole (shown in red in the bottom panels) causes misaligned chromosomes, these misaligned chromosomes satisfy the SAC and result in anaphase lagging and micronuclei formation in daughter cells. See text for details.

### NuMA-mediated k-fiber minus-end clustering is required for proper chromosome segregation

In this study, we found that depletion of the spindle pole-localizing protein NuMA causes chromosome misalignment due to detachment of k-fiber minus-ends from spindle poles in metaphase human cells. Although the majority of k-fiber minus-ends are still clustered, some k-fiber minus-ends are dissociated from one pole as a single or clustered mini spindle-pole (Figure 1D, 2A, S2F, 3A and 4C bottom). Importantly, misaligned chromosomes generated by the k-fiber minus-end detachment maintain chromosome biorientation, and thus are not sensed by the SAC resulting in lagging chromosomes in anaphase and micronuclei formation in interphase (Figure 4C). Our results are consistent with the previous finding that Mad2 does not accumulate on kinetochores after laser ablation of one of the k-fiber pair in metaphase^23^. In addition, inter-kinetochore distance of the misaligned sister kinetochores may be maintained by bridging fibers linking sister k-fibers^24^.

Prior studies established that merotelic kinetochore attachment, in which a single kinetochore is connected to microtubules emanating from different spindle poles (Figure 4C), escapes mitotic checkpoint and causes lagging chromosomes in mammalian cells^25^. Since the merotelic attachment is an abnormal attachment between kinetochore and microtubule plus-ends, our study provides a new concept that detachment of k-fiber minus-ends from focused spindle poles can also cause micronuclei formation by evading mitotic checkpoint mechanisms in mammalian cells (Figure 4C).

Then, how does NuMA maintain k-fiber minus-end clustering? Since NuMA has a C-terminal microtubule-binding domain, which is dimerized by its central coiled-coil^12^, NuMA itself may crosslink focused microtubule pairs^26^. In addition, NuMA recruits the microtubule-binding dynein/dynactin complex to the minus-ends of microtubules in mitosis through its N-terminal region^11^. The resulting dynein-dynactin-NuMA (DDN) complex may concertedly crosslink two relatively separated microtubules using its separated microtubule-binding modules, which may especially be important for centrosome clustering in tetraploid or extra-centrosome containing cells. On the other hand, DDN can also localize to the mitotic cell cortex where it captures astral microtubules to control spindle positioning^15^. Cortical DDN may function for centrosome clustering in cooperation with pole-localized DDN in tetraploid cells, as reported recently^27^. Interestingly, NuMA is over-expressed in some cancer cells and elevated NuMA expression promotes the formation multipolar spindles in cancer cells with extra centrosomes^28^. Considering this and our result, NuMA expression level would need to be tightly regulated to ensure proper bipolar spindle assembly.

In conclusion, our study reveals that maintenance of k-fiber minus-end clustering is critical for proper chromosome segregation. Recent studies indicate that spindle-pole focusing mechanisms are context-dependent^7^ and some pole-focusing factors including ASPM and CDK5RAP2^29^ are related to microcephaly, a disease associated with reduced brain size^30^. It would be important to study whether dysfunction of NuMA and other pole-focusing factors causes chromosome mis-segregation in specific contexts, such as neuronal precursor divisions, for a better understanding of the context-dependent mechanisms and roles of spindle-pole focusing.

## Acknowledgments

We thank Masato Kanemaki for the reagents for the AID2 system. This work was supported by grants from KAKENHI (17H05002 and 21H02481) of the Japan Society for Promotion of Science (JSPS) and the Okinawa Institute of Science and Technology Graduate University, Japan.

## Author contributions

Conceptualization, TK; Investigation, MvT, AG and TK; Formal analysis, MvT, AG and TK; Resources, MvT, AG, SB, and TK; Methodology, MvT and TK; Writing, MvT and TK; Supervision, TK; Funding Acquisition, TK.

## METHODS

### KEY RESOURCES TABLE

#### CONTACT FOR REAGENT AND RESOURCE SHAREING

Further information and requests for resources and reagents should be directed to and will be fulfilled by the Lead Contact, Tomomi Kiyomitsu

#### EXPERIMENTAL MODEL AND SUBJECT DETAILS

Established human tissue culture cell lines, and sequence information about guide RNA and PCR primers used in this study are described in Table S1, S2, and S3, respectively.

#### METHOD DETAILS

- Plasmid Construction
- Cell Culture, Cell Line Generation, and Treatments
- Microscope System and Live Cell Imaging

#### QUANTIFICATION AND STATISTICAL ANALYSIS

- Quantification of Fluorescent intensities and phenotypes
- Statistical Analysis

#### METHOD DETAILS

##### Key resources table

| REAGENT or RESOURCE | SOURCE | IDENTIFIER |
| --- | --- | --- |
| *Chemicals* |  |  |
| SiR-tubulin | Spirochrome | SC002 |
| SiR700-tubulin | Spirochrome | SC014 |
| SNAP Cell 647-SiR | New England BioLabs | S9102S |
| Hoechst 33342 | Sigma-Aldrich | B2261 |
| Nocodazole | Sigma-Aldrich | M1404 |
| RO-3306 | Sigma-Aldrich | SML0569 |
| Puromycin dihydrochloride | Wako Pure Chemical Industries | 160-23151 |
| G-418 solution | Roche | 04727894001 |
| Hygromycin B | Wako Pure Chemical Industries | 084-07681 |
| Blasticidin S hydrochloride | Funakoshi Biotech | CKK-400 |
| Doxycycline hyclate | Sigma-Aldrich | D9891 |
| 3-Indoleacetic acid (IAA) | Wako Pure Chemical Industries | 098-00181 |
| 5-Phenyl-indole-3-acetic acid<br>(5-Ph-IAA) | Kanemaki Lab |  |
| Cytochalasin D | Sigma-Aldrich | D1641 |
| Thymidine | Sigma-Aldrich | T9250 |
| DirectPCR<sup>®</sup> lysis reagent | Viagen Biotech | 302-C |
| *Software and algorithms* |  |  |
| Photoshop, version 23.4.1 | Adobe Inc. | http://www.adobe.com |
| Illustrator, version 26.3.1 | Adobe Inc. | http://www.adobe.com |
| Fiji | <sup>31</sup> | https://fiji.sc/ |
| Metamorph | Molecular Devices | https://www.moleculardevices.com |
| GraphPad Prism 9, version 9.4.1 | GraphPad Software | https://www.graphpad.com |
| Excel | Microsoft Corporation | https://products.office.com/ |

##### Plasmid Construction

Plasmids for CRISPR/Cas9-mediated genome editing and auxin-inducible degron methodology were constructed according to protocols of Natsume et al. ^18^ and Okumura et al., ^15^. To construct donor plasmids containing homology arms for Mis12 (~500-bp homology arms), Mad2 (~200-bp), BubR1 (~200-bp), and TUBG1 (~200-bp), gene synthesis services from Azenta Life Sciences (Suzhou, China) were used. Plasmids and sgRNA sequences used in this study are listed in Supplementary Tables S1 and S2 and will be deposited to Addgene.

##### Cell Culture, Cell Line Generation and Treatments

All HCT116 cell lines were cultured in humidified incubators at 37 °C and 5% CO2 using McCoy’s modified 5A medium (Gibco) supplemented with 10% fetal bovine serum (Gibco) and penicillin-streptomycin (Wako). Knock-in cell lines were generated according to procedures described previously^1^ and screened for mycoplasma contamination using MycoAlert Mycoplasma Detection Kit (Lonza). To activate auxin-inducible degradation, cells were treated with either 2 μg/mL doxycycline (Sigma-Aldrich) and 500 μM indoleacetic acid (IAA, Wako), or 1 μM 5-Phenyl-indole-3-acetic acid (5-Ph-IAA, a kind gift from Masato Kanemaki) for the classical AID^18^ and AID2^19^ systems, respectively. Cells with undetectable signals for the mAID-fusion proteins were analyzed.

Cell lines were generated by transfection with Effectene (Qiagen, Venlo, the Netherlands), followed by selection with the appropriate antibiotics as described previously15. The DNA of single-cell colonies was isolated using DirectPCR lysis solution (Viagen Biotech, Los Angeles, CA), according to the manufacturer’s protocol. Colonies were screened for correct insertion of the fusion cassettes using the primers listed in Supplemental Table S3.

Where indicated, cells were synchronized using 9 μM RO-3306 (Sigma-Aldrich) in culture medium for 20 hours or 2 mM thymidine (Sigma-Aldrich). For NuMA depletion coupled with double thymidine block (Fig 1C and 1D), cells were first arrested in S phase with thymidine for 17 hours. After thymidine washout, cells were grown in fresh medium for 9 hours and then treated with thymidine and Dox for 15 hours. After the 2^nd^ release, cells were incubated with Dox and IAA for ~ 9 hours to deplete NuMA before entering mitosis. Cells were washed with culture medium 3 successive times to release the cells from drug-induced cell cycle arrest. For experiments in Fig 4A-B and S4B-C, cytokinesis was inhibited by addition of 0.2 μM Cytochalasin D (Sigma-Aldrich) to the culture medium for 4 hours after RO-3306 was washed out. For nocodazole experiments, 330 nM nocodazole (Sigma-Aldrich) was added to culture medium 4 hours before imaging and maintained in the medium throughout the imaging session to disrupt microtubule polymerization.

##### Microscope System and Live Cell Imaging

Imaging was performed on a spinning-disc confocal microscope equipped with a 60×1.40 numerical aperture objective lens (Plan Apo λ, Nikon, Tokyo, Japan) and 488, 561, 640 and 685 nm lasers (Coherent, Santa Clara, CA). A CSU-W1 confocal unit (Yokogawa Electric Corporation, Tokyo, Japan) with an ORCA-Flash 4.0 digital CMOS camera (Hamamatsu Photonics, Hamamatsu City, Japan) was connected to an ECLIPSE Ti-E inverted microscope (Nikon) with a perfect focus system for image acquisition. DNA images from cells stained with Hoechst^®^ 33342 (Sigma-Aldrich) were obtained using a SOLA LED light engine (Lumencor, Beaverton, OR).

Live cell confocal imaging was essentially carried out as described before, with minor adjustments. All live cells imaging experiments were carried out in a stage-top incubator (Tokai Hit, Fujinomiya, Japan) maintained at 37° C and 5% CO2. Cells were seeded on glass-bottom dishes (CELLview™, #627860 or #627870, Greiner Bio-One, Kremsmünster, Austria) and pre-stained with 50 nM SiR-tubulin, 50 nM SiR700-tubulin, (all Spirochrome), 120 nM SNAP Cell 647-SiR or 50 ng/mL Hoechst^®^ 33342 (Sigma-Aldrich) for >1 hour before observation where indicated.

For snap shots, five (Figs 1A, S1B and S4B) or fifteen (Fig 4A) z-section images using 1-μm spacing were acquired and maximum intensity projections (Fig 1A and S4B) or single z-section images (Fig S1B) are shown. Signals were linearly adjusted using Fiji and Photoshop to optimize image clarity.

For time-lapse imaging, we either acquired two or three z-section images with 1-μm spacing every 3 or 5 minutes for 2 - 3 hours. Maximum intensity projections (Figs 2A-B, 2E and S2D-I) or single z-section images (Fig 1D) are shown. All time-lapse imaging was performed in the presence of DNA and tubulin dyes as well as with Dox, IAA, 5Ph-IAA and nocodazole where indicated, but of the absence of SNAP substrates.

##### Quantification of fluorescent signals

Fluorescent intensities were quantified using 16-bit images and ImageJ. Mad2 and BubR1 accumulation were determined by measuring the signal intensity of the respective proteins on the kinetochores of interest, and subsequently subtracting the average cellular background signal. Inter-kinetochore distance was measured by drawing and measuring the length a straight line between sister kinetochores in ImageJ.

##### Statistical Analysis

Statistical significance was determined using Graphpad Prism version 9 (GraphPad Software, La Jolla, CA), using two-sided Welch’s t-tests. In Fig 4B and S4C, Z-test (Z-Test Calculator for 2 Population Proportions; https://www.socscistatistics.com/tests/) was used to determine significantly. P values are shown as *: p <0.05, **: p<0.01, ***: p<0.001 and ****: p<0.0001.

## Supplemental Information

**Figure S1.**
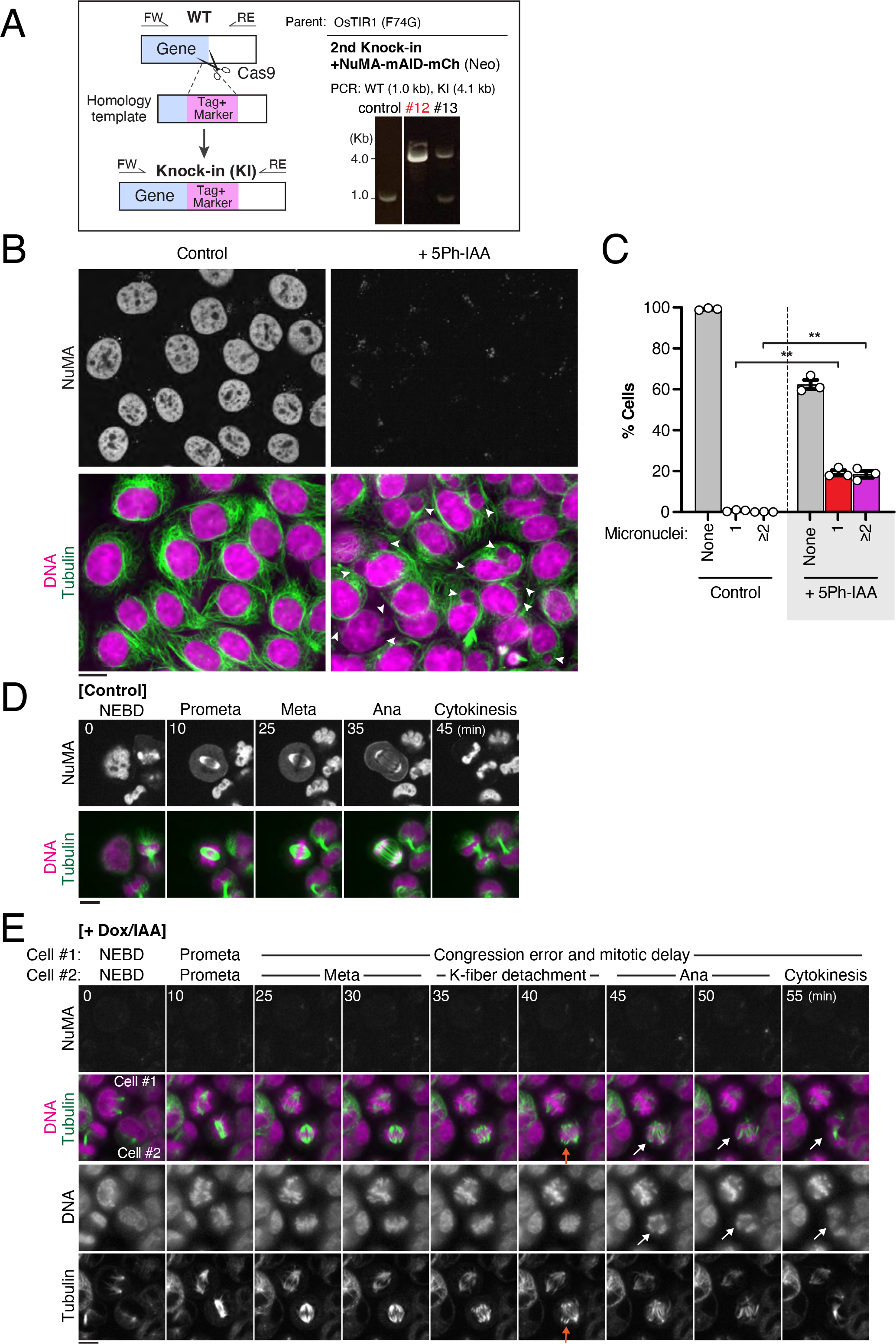
**(A)** Left: Schematic representation of the generation of NuMA-mAID-mCherry knock-in (KI) AID2 cells. The mAID-mCherry cassette, along with neomycin resistant gene (Neo), was inserted at the C-terminus of the *NuMA* loci using CRISPR/Cas9-mediated homology-directed repair. FW: Forward primer, RE: reverse primer. Right: PCR-based genotyping of the *NuMA* gene in control and KI cells that were generated in the AID2 background^19^. A single band of around 4 kb confirms homozygous insertion in the KI clone (#12). **(B)** Representative live-cell fluorescent images showing the presence of micronuclei in interphase NuMA-mAID-mCherry KI cells. Where indicated, endogenous NuMA was degraded by addition of 1 μM 5Ph-IAA 20 hr prior to imaging. Arrowheads indicate micronuclei. **(C)** Quantification of micronuclei formation in control and NuMA-depleted interphase NuMA-mAID-mCherry KI cells. Plotted values are averages of n = 3 independent experiments ± SEM. **(D)** Representative fluorescent images showing time-lapse imaging of mitotic progression in control NuMA-mACF KI cells released from double thymidine block. **(E)** Time-lapse imaging of NuMA-depleted NuMA-mACF KI cells, showing k-fiber minus-end detachments (orange arrow) and anaphase lagging chromosomes and micronucei (white arrows) associated with NuMA deficiency in cell #2. Cell #1 shows a congression defect and consequent mitotic delay. Scale bars = 10 μm.

**Figure S2.**
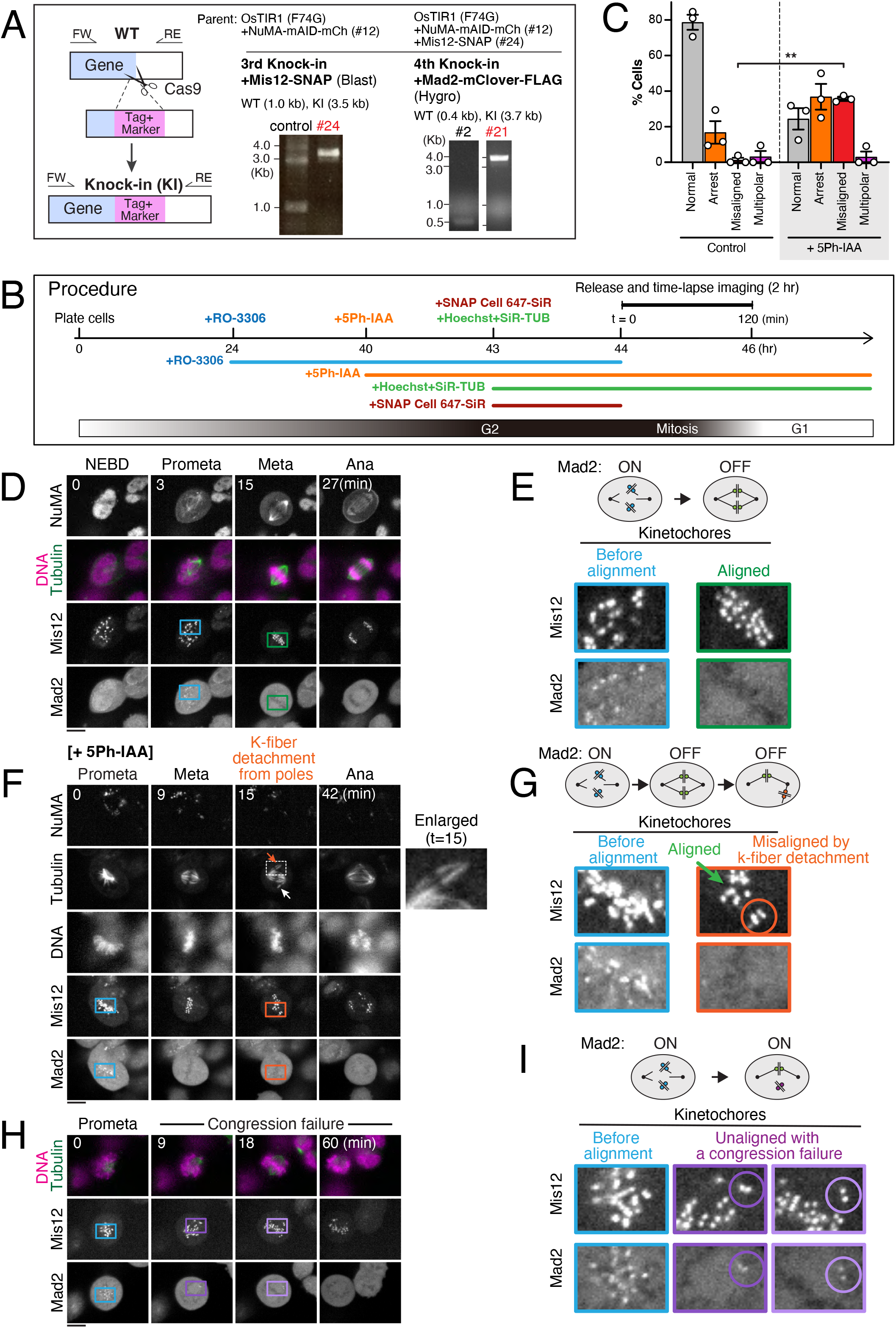
**(A)** Left: Schematic representation of the generation of Mis12-SNAP and Mad2-mClover-FLAG KI cells. The SNAP and mClover cassettes, along with blasticidin S and hygromycin B selection marker genes (depicted as SNAP and mClover for simplicity, respectively), were inserted at the C-terminus of the *Mis12* and *Mad2* loci using CRISPR/Cas9-mediated homology-directed repair. FW: Forward primer, RE: reverse primer. Right: PCR-based genotyping of the *Mis12* and *Mad2* genes in WT and KI cells. A single band of around 3.5 or 3.7 kb confirms homozygous insertion of SNAP or mClover cassette in Mis12 (#24) or Mad2 (#21) gene loci, respectively. **(B)** Outline of the procedure utilized to deplete NuMA in G2 phase followed by live cell imaging of mitotic progression. **(C)** Percentages of control and NuMA-depleted KI cells that either enter anaphase normally (normal), show a prolonged mitotic arrest (arrest), enter anaphase with misaligned chromosomes after detachment of k-fiber minus-ends from focused poles (misaligned) or contain more than 2 spindle poles (multipolar) after release from RO-3306-induced cell cycle arrest. In this condition, ~20% of control cells showed mitotic delay, probably due to the effect of G2 arrest and release using RO-3306. Plotted values are averages of n = 3 independent experiments ± SEM. **(D)** Representative fluorescent images showing time-lapse imaging of mitotic progression in control NuMA-mAID-mCherry KI cells released from RO-3306-induced G2/M block without the addition of 5-Ph-IAA. **(E)** Top: Schematic representation of Mad2 localization at kinetochores before and after alignment on the metaphase plate. Bottom: Enlargement of indicated areas in (D), showing Mad2-positive kinetochores before alignment (e.g. during prometaphase) and Mad2-negative kinetochores after establishment of a focused spindle pole during metaphase. **(F)** Another example of chromosome misalignment and lagging chromosomes generated by NuMA-depletion. As indicated, not only a single (white arrow), but also two focused k-fibers (orange arrow, an enlarged image is shown to the right) are dissociated from the metaphase spindle pole (t = 15), which results in anaphase lagging chromosomes (t = 42). **(G)** Top: Schematic representation of Mad2 localization at kinetochores before, during and after losing alignment on the metaphase plate. Bottom: Enlargement of indicated areas in Figure S2F, showing Mad2-positive kinetochores pre-alignment (e.g. during prometaphase) and Mad2-negative kinetochores after detachment from the focused spindle pole during metaphase. **(H)** Live-cell fluorescent images showing chromosome congression failure in NuMA-depleted (4 hr after addition of 1 μM 5Ph-IAA) KI cells released from RO-3306-induced G2/M arrest. Mad2 localized on misaligned chromosomes with congression defect during the prolonged mitosis. **(I)** Top: Schematic representation of Mad2 localization at kinetochores before alignment, and during failure of chromosome congression. Bottom: Enlargement of indicated areas in Figure S2H, showing Mad2-positive kinetochores before alignment (e.g. during prometaphase) and on unaligned chromosomes (purple circle). Scale bars = 10 μm.

**Figure S3.**
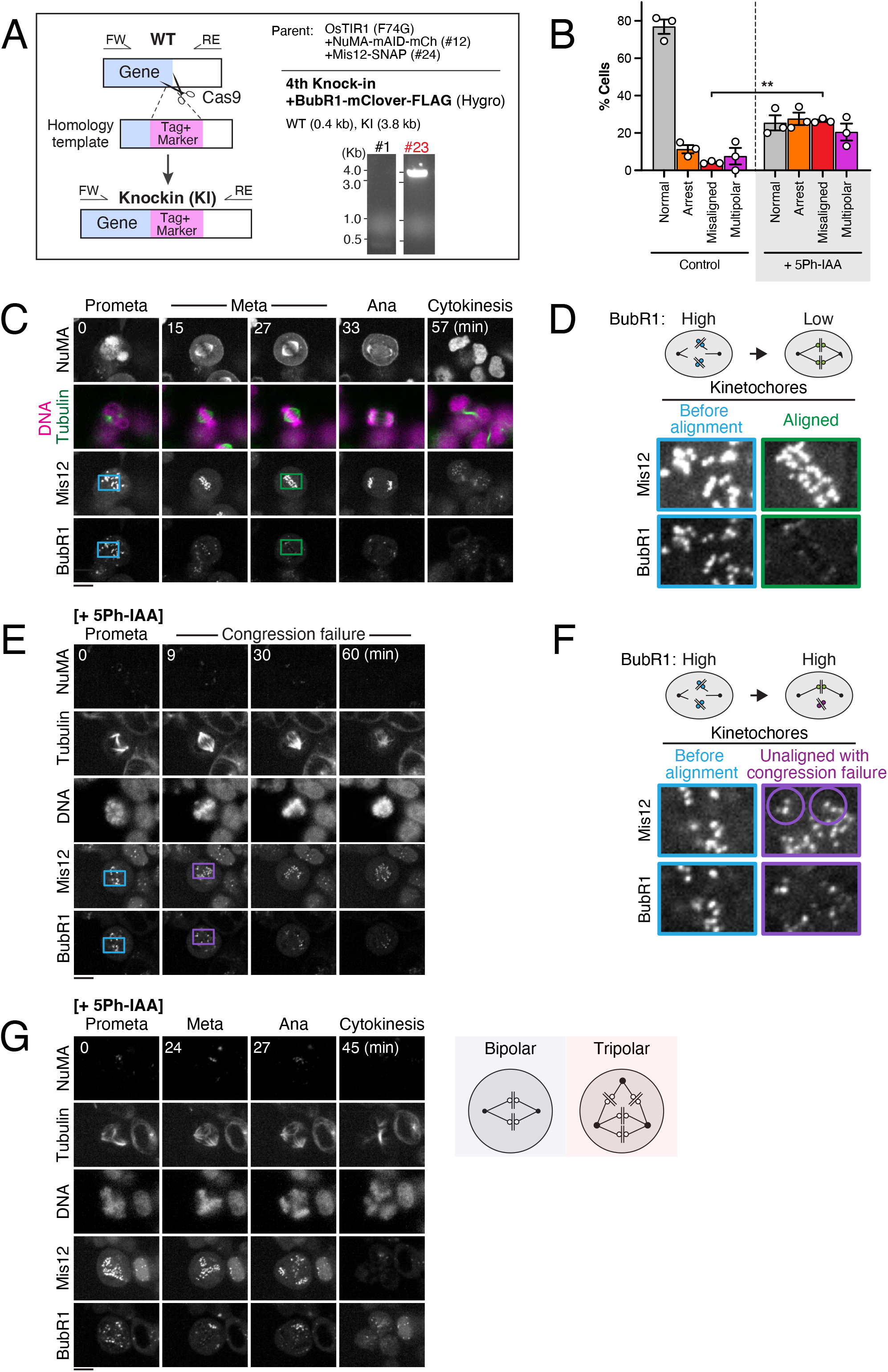
**(A)** Left: schematic representation of the generation of BubR1-mClover KI cells. The mClover cassettes, along with hygromycin B resistant gene (Hygro), were inserted at the C-terminus of the *BubR1* loci using CRISPR/Cas9-mediated homology-directed repair. FW: Forward primer, RE: reverse primer. Bottom: PCR-based genotyping of the *BubR1* genes in WT and KI cells. Right: a single band of 3.8 kb confirms homozygous insertion in the KI clones (#23). **(B)** Percentages of control and NuMA-depleted KI cells that either enter anaphase normally (normal), show a prolonged mitotic arrest (arrest), enter anaphase with misaligned chromosomes after detachment of k-fiber minus-ends from focused poles (misaligned) or contain more than 2 spindle poles (multipolar) after release from RO-3306-induced cell cycle arrest. Plotted values are averages of n = 3 independent experiments ± SEM. **(C)** Representative fluorescent images showing time-lapse imaging of mitotic progression in control cells released from RO-3306-induced G2/M block in the absence of 5-Ph-IAA. **(D)** Top: Schematic representation of BubR1 localization at kinetochores before and after alignment on the metaphase plate. Bottom: Enlargement of indicated areas in (C), showing BubR1 dissociation from aligned kinetochores on metaphase plate. **(E)** Live-cell fluorescent images showing chromosome congression failure and BubR1 accumulation on unaligned chromosomes in NuMA-depleted KI cells released from RO-3306-induced G2/M arrest. **(F)** Top: Schematic overview of BubR1 accumulation on kinetochores before alignment, and during failure of chromosome congression. Bottom: Enlargement of indicated areas in (E), showing BubR1 accumulation on kinetochores before alignment and on unaligned chromosomes (purple circle). **(G)** Top: Schematic representation of bipolar and tripolar (multipolar) spindle. Bottom: Live-cell fluorescent images showing the formation of a tripolar mitotic spindle and tripolar division in NuMA-depleted KI cells released from RO-3306-induced G2/M arrest. Scale bars = 10 μm.

**Figure S4.**
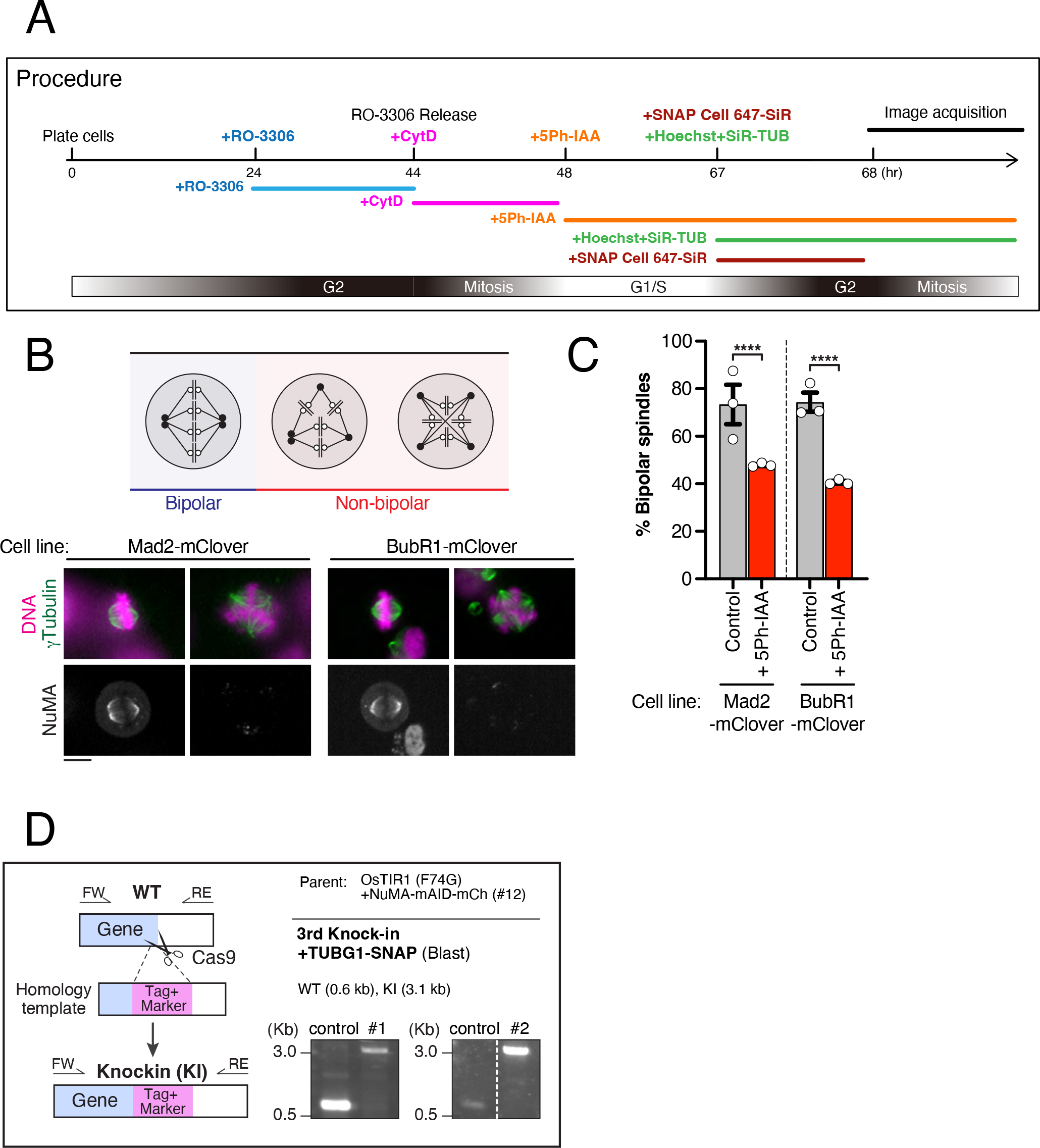
**(A)** Diagram showing the approach to generate tetraploid cells and subsequently determine centrosome clustering in triple (OsTIR1(F74G) / NuMA / TUBG) and quadruple KI (OsTIR1(F74G) / NuMA / Mis12 / Mad2 and OsTIR1(F74G) / NuMA / Mis12 / BubR1) cells. **(B)** Top: Schematic representation of proper (blue, left) and improper (red, right) centrosome clustering in tetraploid cells. Bottom: Representative live-cell fluorescent images of control and NuMA-depleted KI cells showing bipolar and multipolar spindles, marked by SiR-tubulin staining. **(C)** Quantification of the percentage of bipolar spindles in metaphase or early anaphase cells, based on SiR-tubulin staining. Plotted values are average percentages ± SEM from n = 3 independent experiments, in which >20 cells were counted each time. **(D)** Left: Schematic representation of the generation of TUBG1-SNAP KI cells. SNAP cassette, along with the blasticidin S gene (Blast), was inserted at the C-terminus of the *TUBG1* locus using CRISPR/Cas9-mediated homology-directed repair. FW: Forward primer, RE: reverse primer. Right: PCR-based genotyping of the *TUBG1* gene in control and KI cells. A single band of 3.1 kb confirms homozygous insertion in the KI clones.

**Table S1:** Cell lines used in this study.

| No. | Name | Description | Clo<br>ne<br>No. | Plasmids used | Par<br>ent<br>cell | Reference |
| --- | --- | --- | --- | --- | --- | --- |
| 1 | NuMA-mACF | AAVS1::PTRE3G OsTIR1 (Puro), NuMA1::NuMA-mAID-mClover-3FLAG (Neo) | 1 | pTK372 and pTK398 |  | Okumura et al.,<sup>15</sup> |
| 2 | NuMA-mACF + DHC-SNAP | AAVS1::PTRE3G OsTIR1 (Puro), NuMA1::NuMA-mAID-mClover-3FLAG (Neo), DHC1::DHC-SNAP (BSD) | 18 | pTK308 and pTK600 |  | Okumura et al.,<sup>15</sup> |
| 3 | OsTIR1(F74G) | AAVS1::CMV OsTIR1(F74G) (Puro). Puro was removed later. |  |  |  | Yesbolatova et al.,<sup>19</sup> |
| 4 | AID2: NuMA-mAID-mCherry | AAVS1::CMV OsTIR1(F74G), NuMA1::NuMA-mAID-mCherry (Neo) | 12 | pTK372 and pTK989 | 3 | This study |
| 5 | AID2: NuMA-mAID-mCherry +Mis12-SNAP | AAVS1::CMV OsTIR1(F74G), NuMA1::NuMA-mAID-mCherry (Neo), Mis12::Mis12-SNAP (Blast) | 24 | pTK820 and pTK825 | 4 | This study |
| 6 | AID2: NuMA-mAID-mCherry +Mis12-SNAP +Mad2-mClover | AAVS1::CMV OsTIR1(F74G), NuMA1::NuMA-mAID-mCherry (Neo), Mis12::Mis12-SNAP (Blast), Mad2:: Mad2-mClover-FLAG (Hygro) | 21 | pTK1003 and pTK608 | 5 | This study |
| 7 | AID2: NuMA-mAID-mCherry +Mis12-SNAP +BubR1-mClover | AAVS1::CMV OsTIR1(F74G), NuMA1::NuMA-mAID-mCherry (Neo), Mis12::Mis12-SNAP (Blast), BubR1:: BubR1-mClover-FLAG (Hygro) | 23 | pTK875 and pTK1004 | 5 | This study |
| 8 | AID2: NuMA-mAID-mCherry +TUBG1-SNAP | AAVS1::CMV OsTIR1(F74G), NuMA1::NuMA-mAID-mCherry (Neo), TUBG1::TUBG1-SNAP (Blast) | 1 & 2 | pTK611 and pTK624 | 4 | This study |

**Table S2:**
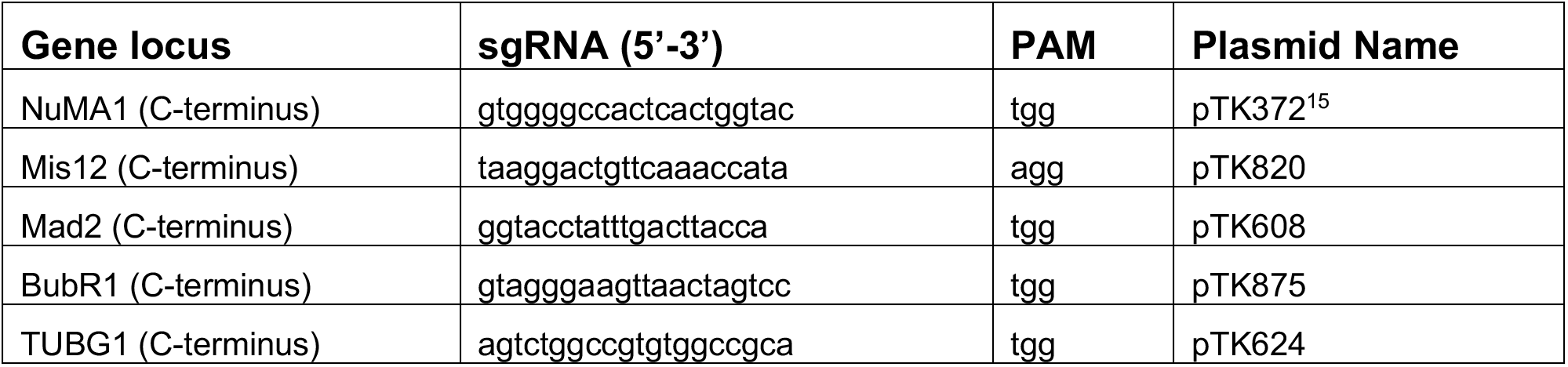
sgRNA sequences for CRISPR/Cas9-mediated genome editing

**Table S3:**
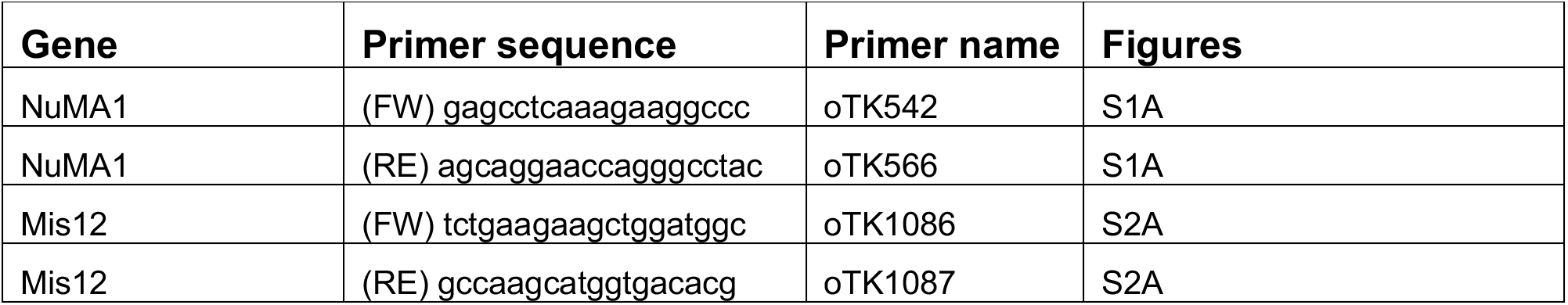
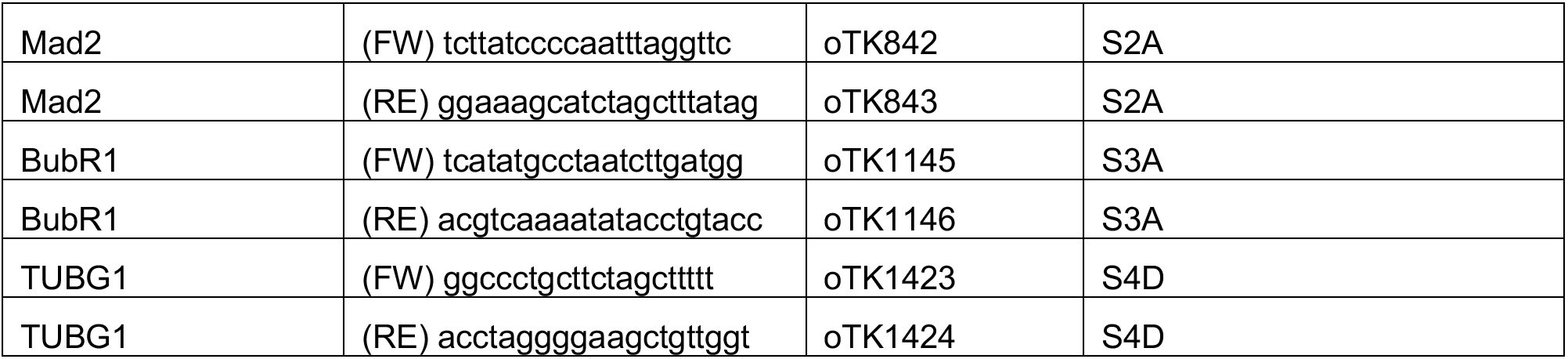
PCR primers used to confirm gene editing

